# Genotype-Specific Electroretinogram Signatures in *Drosophila*: Implications for Neurodegeneration Models Using *w^1118^* Backgrounds

**DOI:** 10.1101/2024.11.29.626125

**Authors:** Nikhil Manuel, Malav Mallipudi, Arya Gajwani, Sravan Gopalkrishna Shetty Sreenivasa Murthy, Daniel C. Jupiter, Balaji Krishnan

## Abstract

*Drosophila melanogaster* serves as a powerful model for studying neurodegenerative diseases, often employing the GAL4-UAS system for targeted gene expression. Electroretinograms (ERGs) provide a robust *in vivo* functional readout of neuronal integrity and are increasingly used to assess disease progression and therapeutic interventions in these models. However, the genetic background upon which these models are built, particularly the widely used *w^1118^* white-eyed mutant, can significantly influence baseline ERG characteristics. This study systematically characterizes ERG responses in wild-type Canton S (CS), *w^1118^*, and a *w^1118^*line carrying a UAS-hPLD1 construct (which includes a mini-white gene). We demonstrate profound differences in ERG amplitudes, waveforms, and responses to varying light stimuli (intensity and duration) between these genotypes, as well as significant sex-specific variations. Notably, *w^1118^* flies exhibit markedly larger ERG amplitudes compared to CS, while the hPLD1 line shows partial compensation. We also introduce a novel quadrant-based analysis of the receptor potential, revealing distinct “fingerprints” for each genotype. These findings underscore that the *w^1118^* background is not electrophysiologically neutral and can intrinsically alter neuronal responses. This has critical implications for interpreting ERG data from neurodegeneration models, as these background effects could mask or mimic disease-related changes. Researchers must consider these baseline differences and potential sex-specific effects to accurately attribute observed ERG phenotypes to the gene or condition under investigation.

**Graphical Abstract:** 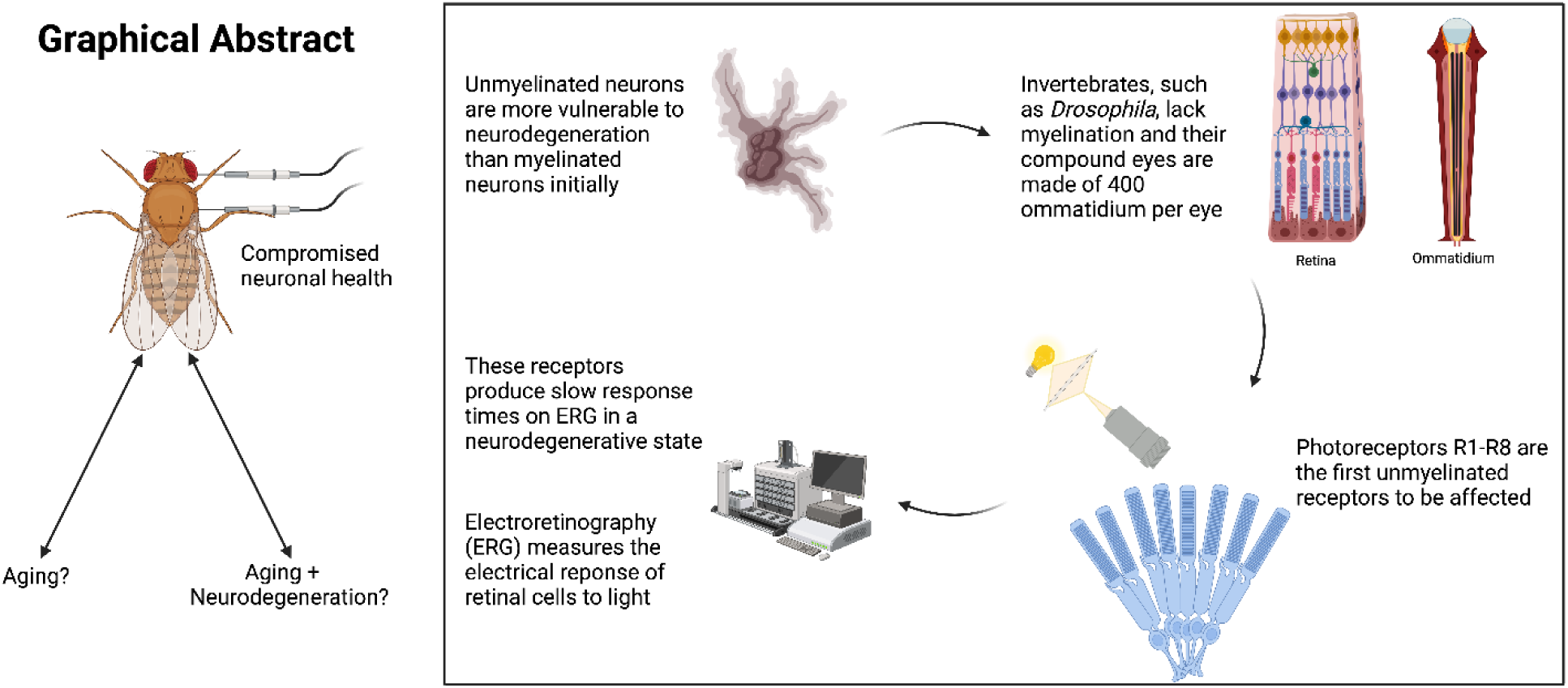

**Highlights:**

- ERG profiles differ significantly between CS, *w^1118^*, and *w^1118^*; UAS-hPLD1 flies.
- The *w^1118^* background, common in GAL4-UAS studies, exhibits distinct ERG features.
- Sex-specific differences in ERG responses are prominent and genotype-dependent.
- UAS-hPLD1 insertion (with mini-white) partially alters the *w^1118^* ERG phenotype.
- Results caution the interpretation of ERG data in *w^1118^* neurodegeneration models.

## Introduction

Neurodegenerative diseases/disorders (NDD) like Alzheimer’s disease (AD) and Parkinson’s disease (PD) pose a growing societal burden, necessitating effective model systems to unravel their complex mechanisms and test potential therapies(‘2025 Alzheimer’s disease facts and figures’). *Drosophila melanogaster* has emerged as a cornerstone in this research due to its sophisticated genetic toolkit, short lifespan, and conserved fundamental biological pathways(Bellen, Tong and Tsuda, 2010). The GAL4-UAS system, in particular, allows for precise spatiotemporal control of transgene expression, enabling researchers to model human diseases by expressing relevant genes (e.g., Tau, Aβ, α-Synuclein) in specific neuronal populations(Caygill and Brand, 2016).

Assessing neuronal function *in vivo* is crucial for understanding disease progression. The *Drosophila* electroretinogram (ERG) offers a powerful, non-invasive (to the neuron) method to measure the summed electrical response of photoreceptor and downstream neurons to light stimuli(Hotta and Benzer, 1969; Belusic, 2011; Krishnan and Wairkar, 2018). It serves as an excellent functional readout for neuronal health and has been successfully employed to study synaptic transmission, ion channel function, and, increasingly, neurodegeneration and aging(Cosens and Manning, 1969; Dolph, Nair and Raghu, 2011). ERGs can detect subtle changes in neuronal activity, potentially even before overt cell death occurs, making them ideal for tracking progressive neurodegeneration(Raghu *et al*., 2009; Thakur *et al*., 2016).

A common practice in *Drosophila* neurodegeneration research is the use of white-eyed mutant backgrounds, most notably *w^1118^*, to facilitate genetic crosses and often because many UAS or GAL4 lines are maintained in this background(Wittmann *et al*., 2001; Muqit and Feany, 2002; Ghosh and Feany, 2004; Ugur, Chen and Bellen, 2016; Şentürk and Bellen, 2018; Ma *et al*., 2022). White-eyed flies lack screening pigments, which simplifies ERG recordings and can lead to larger signals but also alters visual physiology(Pak, Grossfield and White, 1969; Vilinsky and Johnson, 2012). While convenient, the assumption that *w^1118^* is simply a ‘neutral’ background for studying neuronal function, especially in the context of neurodegeneration, has been questioned.

Studies have shown that *white* mutations can inherently lead to retinal degeneration, impact stress resistance, and shorten lifespan. Furthermore, the GAL4 system itself, especially in aging flies, can exhibit changing expression patterns and even induce toxicity, adding another layer of complexity.

Despite the widespread use of ERGs and *w^1118^* flies in neurodegeneration research, a systematic comparison of ERG characteristics between wild-type (CS), *w^1118^*, and *w^1118^*-based transgenic lines (often containing a mini-white rescue marker) is lacking. Such information is critical, as baseline differences could confound the interpretation of results attributed solely to the human gene being studied. This study aims to fill this gap by providing a detailed ERG characterization of CS, *w^1118^*, and a *w^1118^* line expressing human Phospholipase D1 (*w^1118^ and UAS-hPLD1*), which incorporates a mini-white gene(Panda *et al*., 2018; Kankel *et al*., 2020). We demonstrate significant, quantifiable differences between these genotypes and sexes, highlighting potential pitfalls and emphasizing the necessity of considering genetic background effects when using ERGs as a functional measure of neuronal integrity in *Drosophila* neurodegeneration models.

## Results

We systematically recorded and analyzed ERGs from three *Drosophila* genotypes: Canton S (CS) as a wild-type control, *w^1118^* (a common, white-eyed background strain), and *w^1118^;+;UAS-hPLD1* (representing a transgenic line with a partially rescued eye color due to the mini-white gene). We focused on key ERG components: the ‘on-transient’ (initial positive peak), the ‘off-transient’ (positive peak upon light cessation), and the Receptor Potential Amplitude (RPA - the sustained negative potential during illumination). We investigated how these components varied with genotype, sex, light intensity, and stimulus duration.

### 1. Genotype-Specific ERG Waveforms

The fundamental shape and amplitude of the ERG trace differed markedly between genotypes (**Figure 1**). CS flies exhibited a characteristic ERG with clear on- and off-transients and a moderate RPA. In contrast, *w^1118^* flies showed significantly larger RPAs and often reduced or altered transient components, consistent with previous observations in white-eyed mutants. The *w^1118^;+;UAS-hPLD1* line, with its partially restored pigmentation, displayed an intermediate phenotype, often with bimodal characteristics during the receptor potential, especially at longer stimulus durations. These basic differences highlight that the genetic background profoundly influences the ERG response.

**Figure 1.**
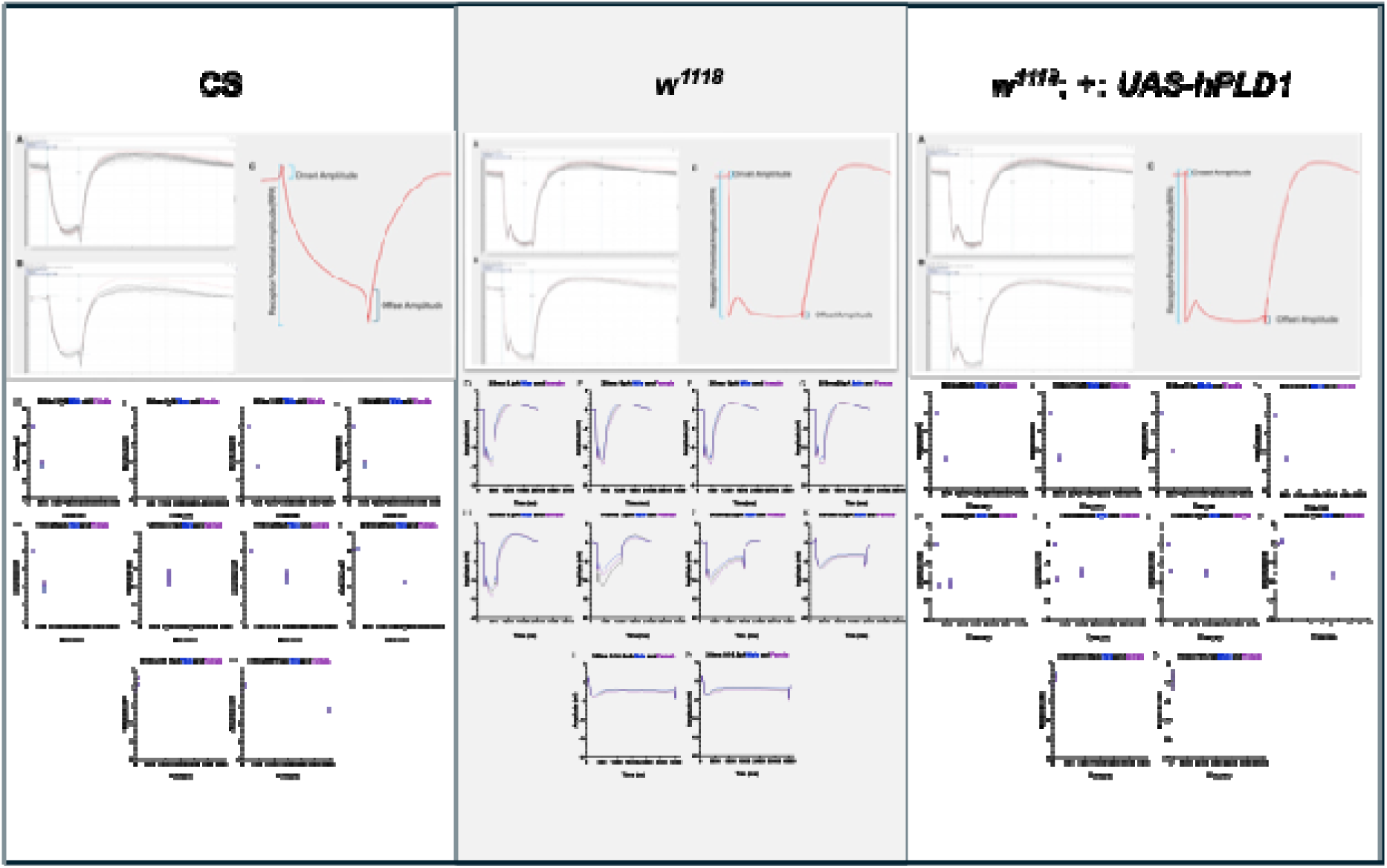
Electroretinogram (ERG) recordings from three *Drosophila melanogaster* genotypes: wild-type Canton S (CS, left column), white-eyed mutant (*w^1118^*, middle column), and a transgenic line expressing human Phospholipase D1 (hPLD1) cDNA downstream of a UAS sequence in the *w^1118^* background (*w^1118^;+;UAS-hPLD1*, right column). Panel A represents the raw ERG trace including electrical interference. **Panel B** represents the ERG trace after filtration of electrical interference, resulting in a smoothened waveform using the Clampfit software. **Panel C** provides an overall hand-drawn representation highlighting key characteristics of the ERG trace, including the Onset Amplitude, Offset Amplitude, and Receptor Potential Amplitude (RPA). **Panels D-M** (distributed across the three genotypes as indicated by their column placement relative to A-C) illustrate ERG responses under varying stimulation conditions. **Top panels within D-G and comparable positions in other genotype columns** display ERG traces elicited by different light stimulation intensities (e.g., specific pA values if discernible, though not explicitly labeled with values in the image) all delivered for a 350ms duration. **Middle panels within H-K and comparable positions in other genotype columns** illustrate ERG patterns resulting from increasing durations of light stimulation (e.g., 500ms, 1000ms, 1500ms, 2000ms as suggested by axis labels) at a consistent, representative intensity. These traces highlight differences in onset, offset, and receptor potential amplitude as a function of stimulus duration. **Lower panels within L-M and comparable positions in other genotype columns s**how ERG patterns in response to either incrementally increasing (e.g., 5.5 pA to 9.5 pA) or decreasing (e.g., 9.5 pA to 5.5 pA) light intensities, each for a 350ms duration, demonstrating the dynamic response of the retina to varying light levels. While this hPLD1 construct is not active without a specific GAL4 driver, these recordings explore baseline similarities and differences in ERG characteristics when this putative genetic tool (intended for studying human neurological conditions) is present.

### 2. Impact of Light Intensity on ERG Components

We examined responses to increasing light intensities (5 pA to 20 pA) at a constant duration (300 ms, based on original Figure 12 legend, although other intensities/durations were tested). Across all genotypes, increasing intensity generally led to larger response amplitudes. However, significant genotype- and sex-specific differences were observed (**Figure 2**).

- **On-Transient:** CS and *w^1118^;+;UAS-hPLD1* groups showed significant differences between sexes, whereas *w^1118^* males and females responded almost identically.
- **RPA:** *w^1118^* and *w^1118^;+;UAS-hPLD1* groups showed significant differences between sexes, while CS did not. The *w^1118^* genotype consistently showed the largest RPA.
- **Off-Transient:** Both *w^1118^*and *w^1118^;+;UAS-hPLD1* groups showed significant sex differences at all intensities, unlike CS.

**Figure 2:**
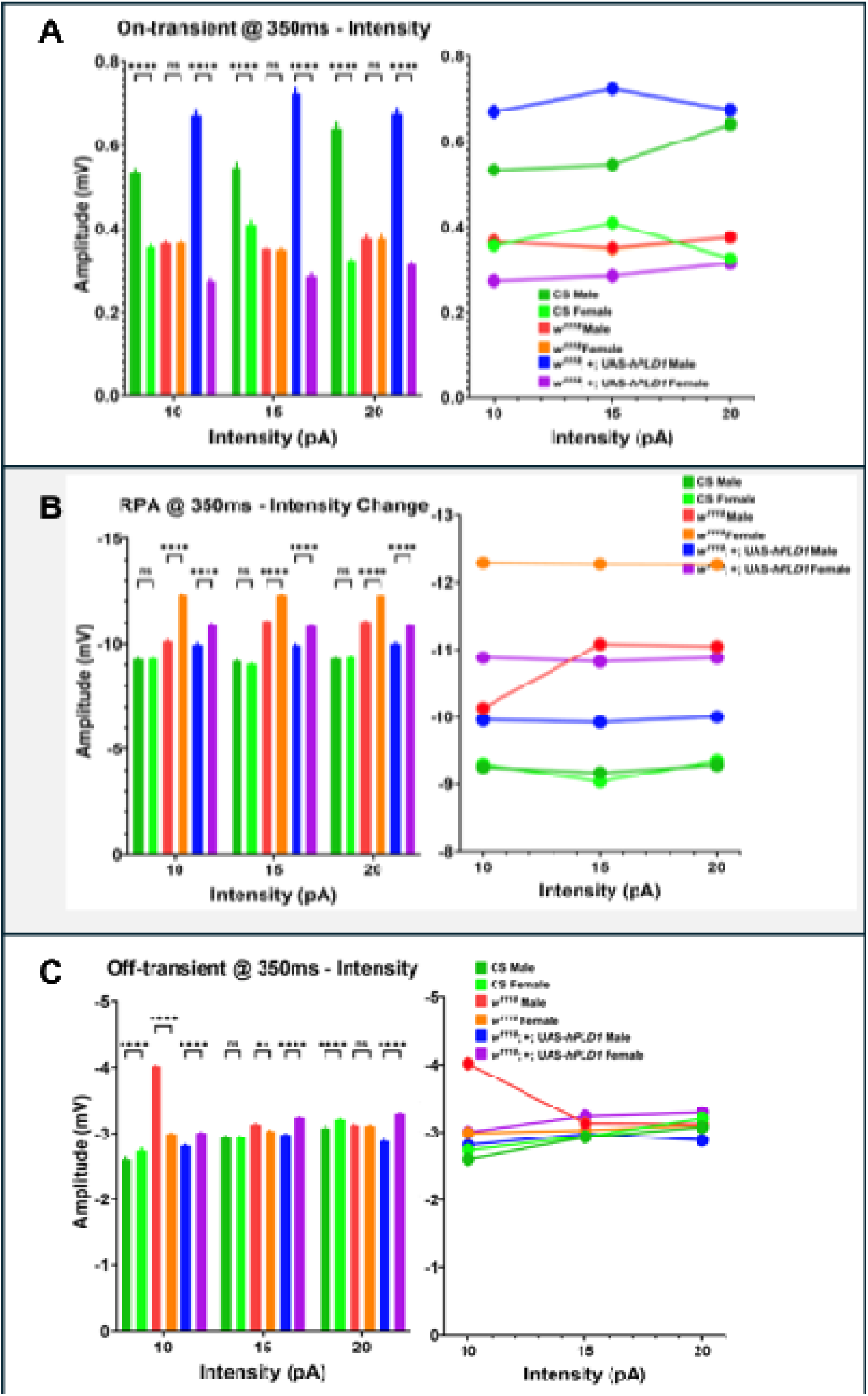
Sex-specific differences in electroretinogram (ERG) parameters across different light intensities in control (CS), mutant (*w^1118^*), and transgenic *Drosophila melanogaster*. All measurements were taken with a light stimulus duration of 350ms. Bar graphs (left within each panel) show mean ± SEM with statistical significance indicated between sexes within each genotype and intensity level. Corresponding line graphs (right within each panel) illustrate these trends. Genotypes and sexes are color-coded as follows: CS Male (light green), CS Female (dark green), *w^1118^* Male (orange), *w^1118^* Female (red), *UAS-hPLD1* Male (blue), and *UAS-hPLD1* Female (purple). **A On-transient Amplitude @ 350ms – Intensity -** At all tested light intensities (10, 15, and 20 pA), on-transient responses showed significant sex differences (****p < 0.0001) in the control CS group and the *UAS-hPLD1* group. Conversely, the *w^1118^* mutant group sexes exhibited no significant differences (ns), with nearly identical and superimposed responses on the line graph (right). **B Receptor Potential Amplitude (RPA) @ 350ms - Intensity Change -** Across all tested intensities, RPA responses were significantly different between sexes (****p < 0.0001) for both the *w^1118^* mutant genotype and the *UAS-hPLD1* genotype. In contrast, the control CS group did not present statistically significant changes (ns) between sexes in RPA. **C Off-transient Amplitude @ 350ms – Intensity -** The off-transient response was statistically different between sexes at all tested intensity levels for both the *w^1118^* mutant group (****p < 0.0001) and the *UAS-hPLD1* group (**p < 0.01). The control CS group, however, did not show significant sex differences (ns) in their off-transient responses. For all panels, statistical analysis was performed using a repeated measures (mixed-effects model) 2-way ANOVA, followed by Sidak’s multiple comparisons test with a single pooled variance. Significance is denoted as ns (not significant), ** (p < 0.01), or **** (p < 0.0001).

These results indicate that both genotype and sex interact to determine the response to varying light strengths, emphasizing the need to analyze these factors separately.

### 3. Impact of Stimulus Duration on ERG Components

We also tested the effect of increasing stimulus duration (300 ms to 2000 ms) at a constant intensity (5.5 pA). Again, complex interactions emerged (**Figure 3**).

- **On-Transient:** Significant sex differences were observed in both CS and *w^1118^;+;UAS-hPLD1* groups at all intervals, with the hPLD1 group showing larger differences. The *w^1118^*group showed no significant sex difference.
- **RPA:** The *w^1118^;+;UAS-hPLD1* group showed sex differences only at the longest interval (2000 ms). *w^1118^*generally maintained the largest RPA.
- **Off-Transient:** Differences between sexes became more pronounced in *all* groups as the stimulus duration increased.

**Figure 3:**
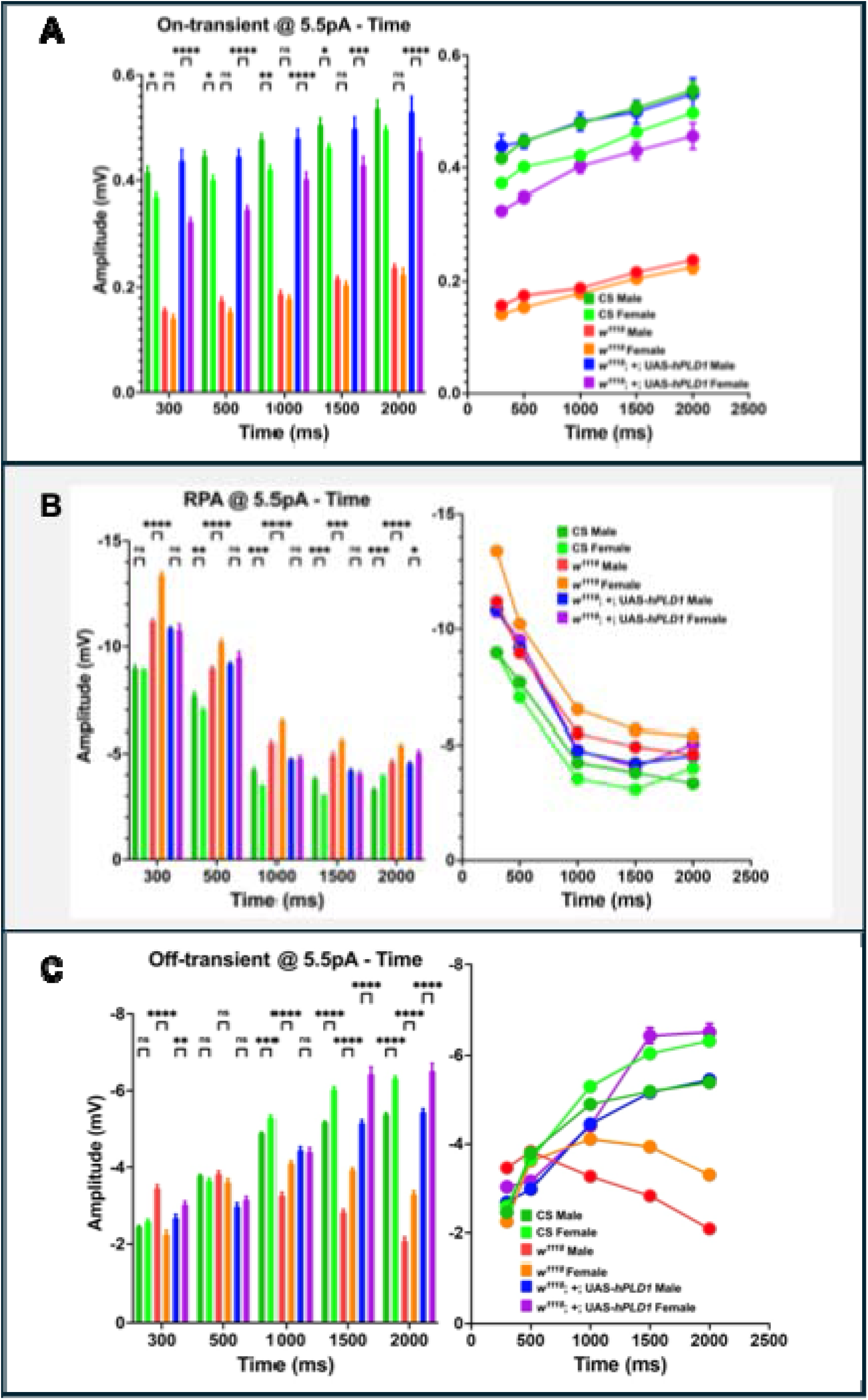
Sex-specific differences in electroretinogram (ERG) parameters as a function of increasing light stimulus duration (300, 500, 1000, 1500, and 2000 ms) at a constant intensity of 5.5pA in control (CS), mutant (*w^1118^*), and transgenic *Drosophila melanogaster*. Responses are shown for control CS (males light green, females dark green), mutant *w^1118^* (males orange, females red), and *UAS-hPLD1* (males blue, females purple). Bar graphs (left within each panel) show mean ± SEM with statistical significance between sexes indicated. Line graphs (right within each panel) illustrate overall trends. **A On-transient Amplitude @ 5.5pA –** Time - As the stimulus duration increased with constant intensity, the on-transient response was significantly different between sexes in both the control CS group (significant at all intervals: 300ms, 500ms, 1000ms, 2000ms with ****p < 0.0001; 1500ms with ***p <; 0.001) and the *UAS-hPLD1* group (****p < 0.0001 at all intervals). The differences in the *UAS-hPLD1* group were particularly pronounced. In contrast, the *w^1118^* mutant group did not show significant sex differences (ns) in on-transient amplitude at any of the tested time intervals. **B RPA @ 5.5pA – Time -** As the stimulus duration increased with constant intensity, the Receptor Potential Amplitude (RPA) in the *UAS-hPLD1*group was not significantly different (ns) between sexes at shorter intervals (300, 500, 1000 ms) but demonstrated significant differences at longer intervals (1500 ms: *p < 0.05; 2000 ms: ***p < 0.001). The *w^1118^* mutant group showed a significant sex difference (*p < 0.05) only at the 300 ms interval, with no significant differences (ns) at other durations. The control CS group did not exhibit significant sex-based differences (ns) in RPA at any tested interval. **C Off-transient @ 5.5pA – Time -** As the stimulus duration increased with constant intensity, the off-transient response in the *w^1118^* mutant group was consistently and highly significantly different (****p < 0.0001) between sexes at all tested time intervals. For the *UAS-hPLD1* group, sex differences in the off-transient response were not significant (ns) at shorter durations (300, 500 ms) but emerged and increased in significance at longer durations (1000 ms: *p < 0.05; 1500 ms: **p < 0.01; 2000 ms: ****p < 0.0001). The control CS group showed no significant sex differences (ns) in off-transient amplitude at any interval. For all panels, statistical analysis of sex differences within each genotype at each time point was performed using a repeated measures (mixed-effects model) 2-way ANOVA, followed by Sidak’s multiple comparisons test with a single pooled variance. Significance is denoted as ns (not significant), * (p < 0.05), ** (p < 0.01), *** (p < 0.001), or **** (p < 0.0001).

This suggests that the temporal processing of visual information is also heavily influenced by genotype and sex.

### 4. Fine-Scale Intensity Responses

To assess sensitivity to smaller changes, we tested responses to intensities between 5.5 pA and 9.5 pA at a constant 350 ms duration. Even within this smaller range, significant sex differences were observed across *all* ERG components and in *all* groups (**Figure 4**). This underscores the pervasive nature of sex-based differences in visual processing, even at relatively low and similar light levels, and reinforces the importance of accounting for sex in ERG studies.

**Figure 4:**
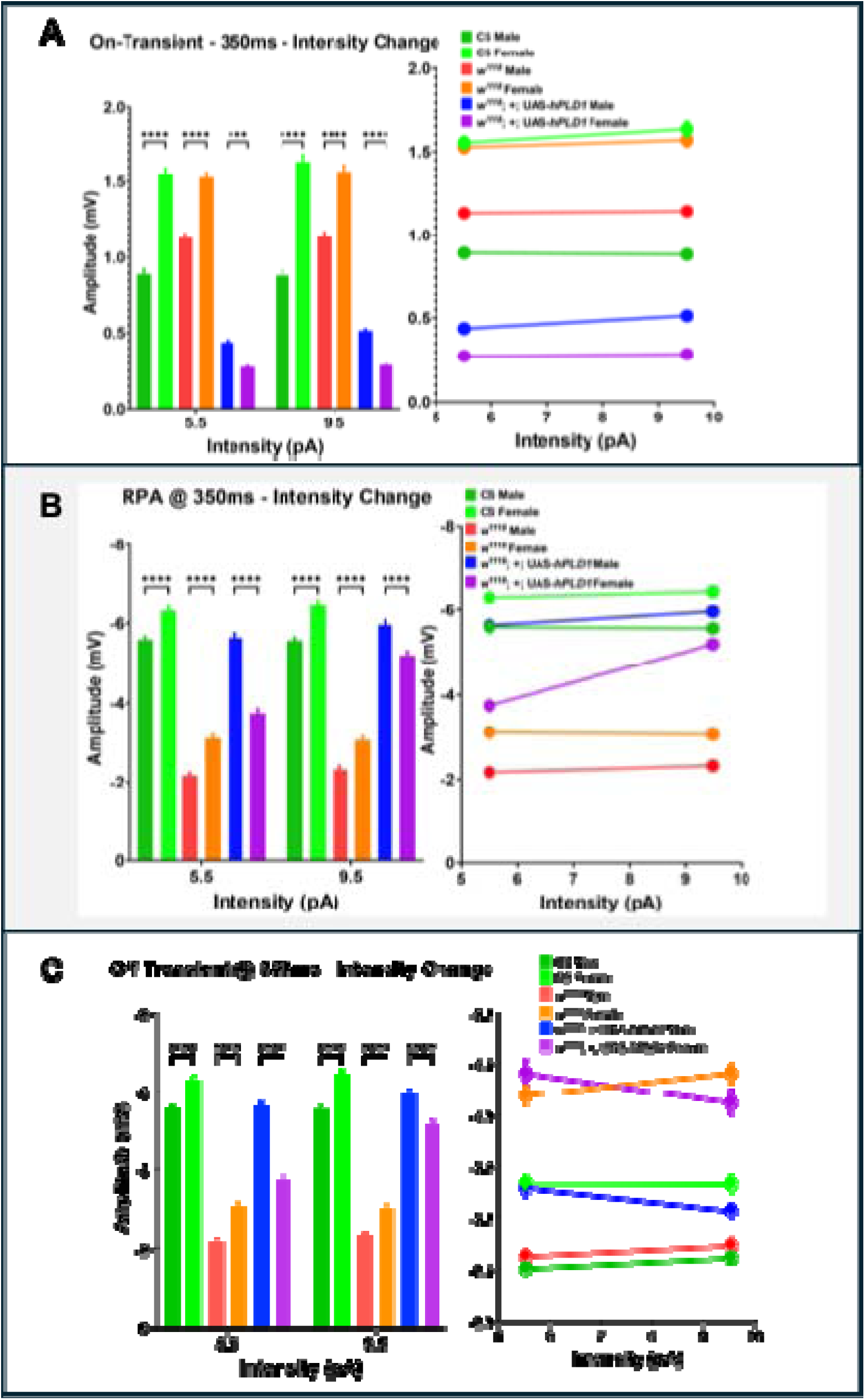
Sex-specific differences in electroretinogram (ERG) parameters as a function of increasing light intensity (5.5 and 9.5 pA) at a constant stimulus duration of 350ms in control (CS), mutant (*w^1118^*), and transgenic *Drosophila melanogaster*. Responses are shown for control CS, mutant *w^1118^*, and transgenic *UAS-hPLD1*. Bar graphs (left within each panel) show mean ± SEM with statistical significance between sexes indicated. Line graphs (right within each panel) illustrate overall trends. **A On-Transient - 350ms - Intensity Change** - As the intensity increased with a constant 350ms interval, the trend seen was that the on-transient response showed significant sex differences (****p < 0.0001) in the control CS group and the *UAS-hPLD1* group at all tested intensity levels (5.5 and 9.5 pA). In contrast, the *w^1118^* mutant group did not exhibit significant sex differences (ns) in on-transient amplitude at any of these intensities. **B. RPA @ 350ms - Intensity Change** - As the intensity increased (to a smaller extent than seen in previous figures 11-16) with a constant 350ms interval, the trend seen was that the RPA was significantly different between sexes (****p < 0.0001) in the *w^1118^* mutant group and the *UAS-hPLD1* group at all tested intensity levels. However, the control CS group did not show significant sex-based differences (ns) in RPA at these intensities. **C. Off-Transient @ 350ms - Intensity Change** - As the intensity increased with a constant 350ms interval, the trend seen was that the off-transient response was significantly different between sexes in the *w^1118^* mutant group (****p < 0.0001) at all tested intensity levels. The *UAS-hPLD1* group also showed significant sex differences (**p < 0.01 at 5.5pA and *p < 0.05 at 9.5pA). The control CS group, however, did not show significant sex differences (ns) in off-transient amplitude at these intensities. For all panels, statistical analysis of sex differences within each genotype at each intensity point was performed using a repeated measures (mixed-effects model) 2-way ANOVA, followed by Sidak’s multiple comparisons test with a single pooled variance. Significance is denoted as ns (not significant), * (p < 0.05), ** (p < 0.01), *** (p < 0.001), or **** (p < 0.0001).

### 5. Novel Quadrant Analysis of Receptor Potential Dynamics

To capture more subtle differences in the ERG waveform, especially during the sustained response (RPA), we developed a novel analysis method. We divided the average ERG trace into distinct zones based on its first derivative (**Figure 5 A-C**). We then focused on the “Mid-Impulse” zone (orange), which represents the RPA, and plotted its trajectory in a centered coordinate system (**Figure 5 D-R**). This “quadrant profile” provides a visual fingerprint of how the RPA evolves over time. We observed distinct patterns for each genotype at different stimulus durations. For example, at shorter durations (≤ 500ms), no genotype entered the third quadrant, but differences in traversal through quadrants I, II, and IV were apparent.

**Figure 5:**
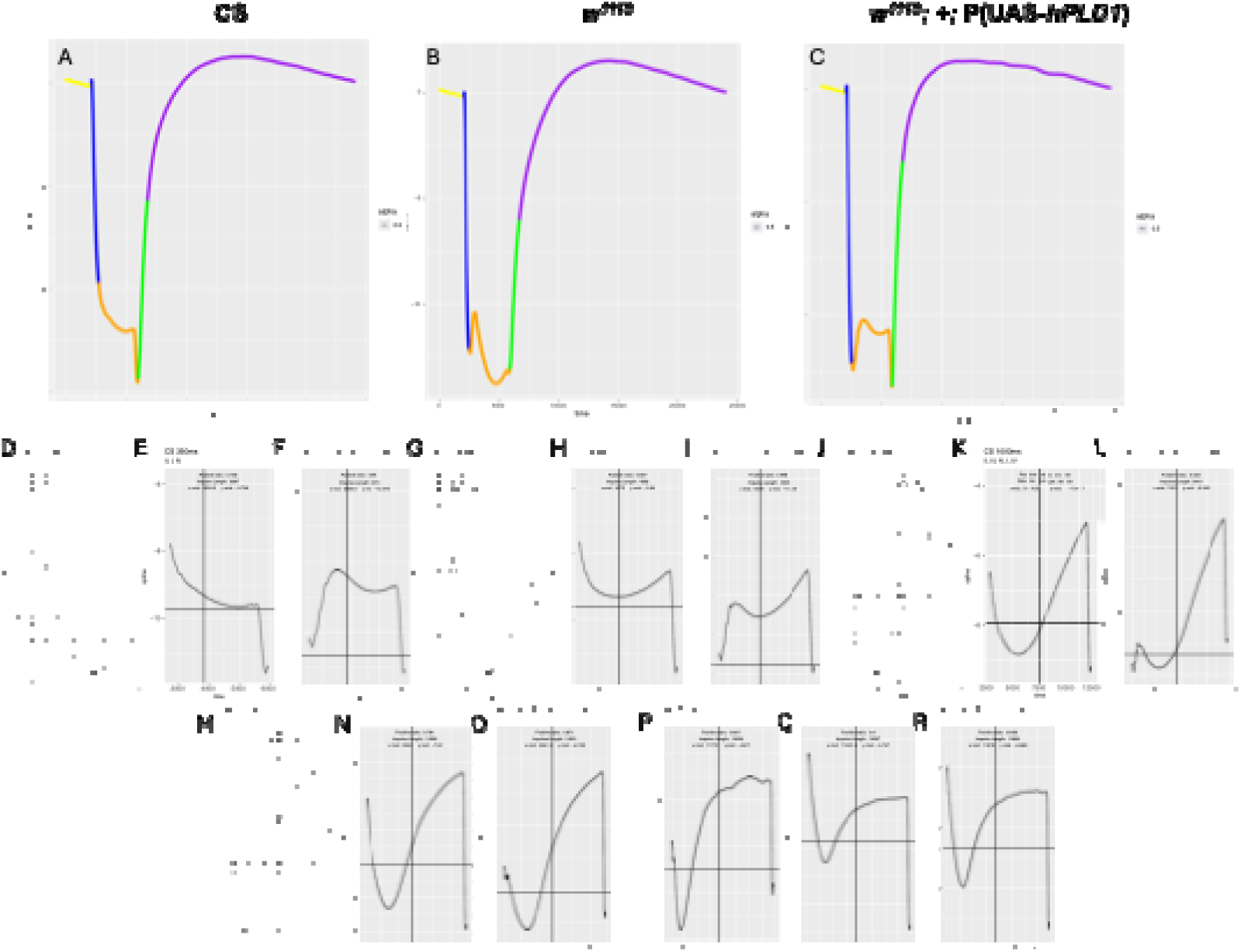
**Segmentation and Detailed Analysis of the Mid-Impulse Phase of ERG Traces in *Drosophila***. Panels A, B, and C display representative average ERG traces for a 350ms pulse of light at 5.5pA, corresponding to genotypes CS, *w^1118^*, and *UAS-hPLD1*, respectively. These traces are segmented into distinct regions based on the first derivative of the signal to highlight specific profile characteristics - **Yellow (Pre-Impulse) r**epresents all data points before the first point where the derivative is less than −0.005. **Blue (Impulse-on-Drop)** includes all points where the derivative is less than −0.005, up to and including the point immediately before the first transition back to a derivative greater than or equal to −0.005. **Orange (Mid-Impulse)** encompasses all points immediately following the first transition from a derivative < −0.005 to >= −0.005, up until right before the first point where the first derivative becomes greater than 0.005. **Green (Impulse Off-Rise)** consists of all points where the first derivative is greater than 0.005, characterizing the initial recovery phase. **Purple (Post-Impulse)** includes all points after the Impulse Off-Rise, representing the later part of the recovery towards baseline. **Panels D through R** provide an expanded view of the **Orange (Mid-Impulse)** portion of ERG traces for the three genotypes across five different stimulus durations (350ms, 500ms, 1000ms, 1500ms, and 2000ms, arranged by column). These detailed views were generated to further elucidate the nature of unique patterns observed within this phase. For these expanded plots: The **x-axis is centered** halfway between the first peak in the On-Drop (if present, otherwise the last peak in the Pre-Impulse) and the last peak occurring before the final transition of the derivative from positive to negative within the Mid-Impulse. The **y-axis is centered** at the halfway point between the y-value at the beginning of the Mid-Impulse and the y-value of the last peak within the Mid-Impulse. The analysis of Cartesian **quadrants (I, II, III, IV) traversed by the traces** in these expanded Mid-Impulse views (indicated in each sub-plot, e.g., “Quadrants: I, IV”) offers insights into the distinct response dynamics of each genotype (further detailed in Table 1). Key observations from this analysis include (i) At stimulation times of 500ms and below, the third quadrant (III) is reportedly never visited by any of the genotypes. (ii) There are notable differences in quadrant traversal patterns between genotypes, particularly in the 500-1500ms stimulus duration zone, compared to responses at 350ms or 2000ms. Furthermore, “positive/negative ratios” (defined as the ratio of the area under the curve in quadrants I and II to the area above the curve in quadrants III and IV) are described as differing substantially at shorter stimulus durations and converging at longer durations.

**Table 1:**
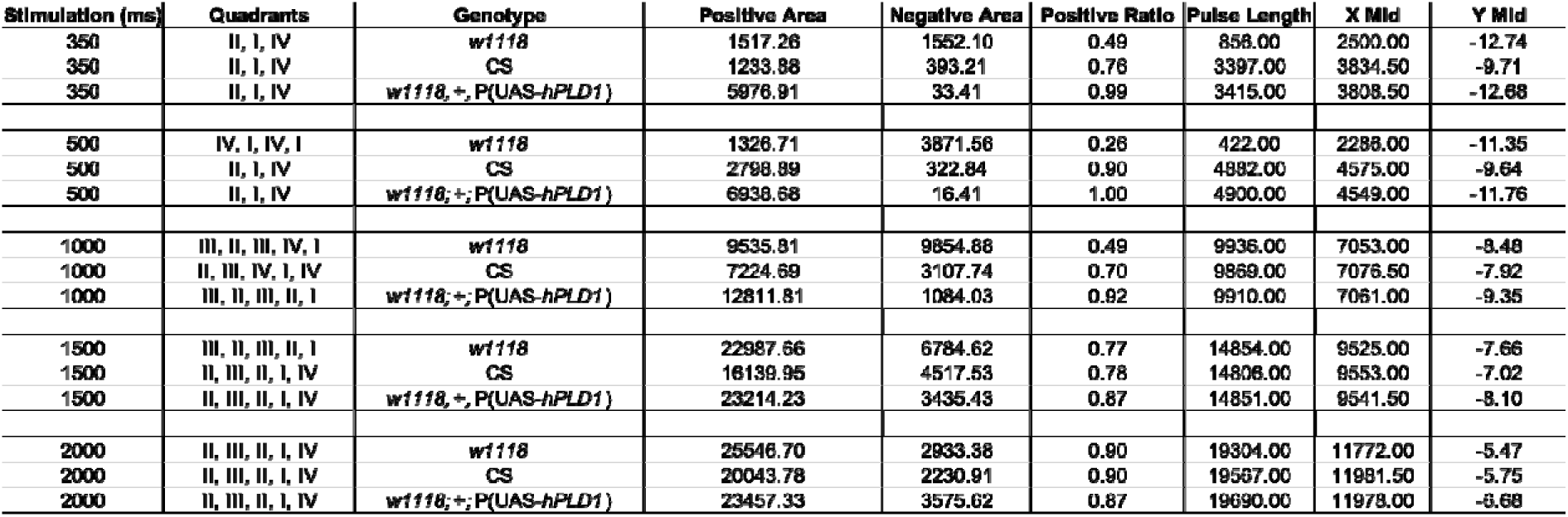
Comparison of the genotypes measured using the quadrant profile associated with the peak of the RPA. When stimulation time (ms) when increased from 350 to 2000ms with a light intensity at 5.5pA, we were able to appreciate the following patterns. Except for 1000ms, the quadrant profile of the human PLD1 insert line parallels with CS, suggesting that the presence of pigmentation affects the RPA nadir characteristics. Given that there is a range of pigmentation reported with the P-element inserts depending on the location, this criteria may provide a quantification to stratify different UAS-lines based on the ERG responses as noted here. The positive to negative quadrants show variable ratios and are genotype dependent but also show correspondence with stimulation lengths suggesting multiple quantifiable metrics to further assess neuronal integrity. Additional parameters that we assessed include the pulse length, midpoint for X-axis and Y-axis that show patterns which can be quantified further.

We quantified these quadrant profiles by noting the sequence of quadrants traversed and calculating the ratio of positive (area under I & II) to negative (area above III & IV) areas. This analysis revealed interesting parallels and divergences (**Table 1**). For instance, the quadrant profile of the hPLD1 line often mirrored the CS line, particularly at durations other than 1000ms, suggesting the partial pigment restoration influences RPA dynamics. The positive-to-negative ratios also showed genotype- and duration-dependent variations. This quantifiable approach offers additional metrics beyond simple amplitude measurements to assess neuronal integrity and differentiate between genotypes.

## Discussion

The primary finding of this study is that *Drosophila* ERG responses are significantly influenced by the genetic background and sex of the fly, with particular relevance for the commonly used *w^1118^* strain. Our results demonstrate that *w^1118^*flies are not electrophysiologically equivalent to wild-type CS flies, exhibiting markedly different ERG amplitudes and waveforms. This confirms and extends previous reports suggesting that the *white* mutation, beyond its eye color phenotype, has physiological consequences, including potential retinal degeneration(Pak, Grossfield and White, 1969; Vilinsky and Johnson, 2012). The larger amplitudes seen in *w^1118^* are likely due to the lack of screening pigments, allowing more light to reach photoreceptors and increasing the collective current generation. However, this lack of pigment also impairs normal light adaptation and can lead to prolonged depolarization under certain conditions, fundamentally altering visual processing.

The *w^1118^;+;UAS-hPLD1* line, carrying a mini-white gene, showed an intermediate phenotype. This suggests that even partial restoration of eye pigmentation can significantly alter ERG responses, bringing them closer to, but not identical with, the CS profile. The bimodal RPA observed in this line might represent a complex interplay between the partially restored pigment-mediated light filtering and the underlying *w^1118^*physiology, or perhaps an effect of the hPLD1 expression itself, although differentiating these requires further study. Regardless, it highlights that even standard components of transgenic constructs (like mini-white) can impact the ERG.

The pervasive sex-specific differences we observed, particularly in the *w^1118^* and hPLD1 lines, are crucial. These differences are not consistent across all genotypes or all ERG parameters, indicating complex genetic and physiological interactions. Failure to account for sex in ERG studies could lead to increased variability and potentially erroneous conclusions, especially when comparing different genotypes or treatment groups.

These findings have significant implications for *Drosophila* neurodegeneration research. Many studies express human disease-associated genes using the GAL4-UAS system in a *w^1118^* background and use ERGs to assess functional decline. Our data strongly suggest that any observed ERG changes in such models must be interpreted with caution. Is a change in ERG amplitude or waveform due to the expressed human gene, or is it an interaction with the inherent *w^1118^* phenotype? For instance, a disease model might show a *decrease* in the large *w^1118^* RPA; this could be interpreted as neurodegeneration, but it might simply reflect a partial ‘rescue’ towards a CS-like phenotype, or an interaction with the *w^1118^*background’s known susceptibility to degeneration. Furthermore, studying aging effects requires extra care, given that ERGs and GAL4 expression both change with age, potentially confounding results.

Our novel quadrant analysis provides a more nuanced way to characterize ERG waveforms, potentially offering a sensitive metric to distinguish subtle changes in neuronal function that might be missed by simple amplitude measurements. The distinct “fingerprints” we observed (**Figure 5**) and their quantification (**Table 1**) could be developed into a valuable tool for stratifying different transgenic lines or assessing the impact of neurodegeneration on signal processing dynamics.

In conclusion, while *Drosophila* ERGs remain a vital tool for neurodegeneration research, our study emphasizes the critical importance of understanding and accounting for the effects of genetic background and sex. The *w^1118^* strain, while widely used, introduces significant baseline differences in ERG responses. Researchers must use appropriate controls (including *w^1118^*without the transgene and potentially CS or other wild-types), analyze sexes separately, and consider the potential interactions between their gene of interest and the background physiology. This study provides crucial baseline data and highlights the need for careful experimental design and interpretation to ensure that ERG assessments accurately reflect the progression of neurodegenerative states in *Drosophila* models.

## Abbreviations

GAL4/UAS: GAL4/Upstream Activating Sequence
AD: Alzheimer’s Disease
BDSC: Bloomington Drosophila Stock Center
APP: Amyloid Precursor Protein
APPL: Amyloid Precursor protein Homolog
MAPT: Microtubule-Associated Protein Tau
BACE1: beta-site amyloid precursor protein cleaving enzyme 1
ERG: Electroretinogram
Elav GAL4: Embryonic Lethal Abnormal Vision GAL4
*w^1118^*: *white1118*
PLD1: Phospholipase D Isoform 1
RPA: Receptor Potential Amplitude
CS: Canton S

## Funding Sources & Ethics Statement

This work was supported by research grants from the NIH R01 AG063945(B.K.), NIH R21 AG059223(B.K.), AARG-17-533363(B.K), The Don and Nancy Mafrige Professor in Neurodegenerative Disease Endowment (B.K.), Mitchell Center for Neurodegenerative Diseases (B.K.).

This study was conducted in accordance and compliance with the National Institutes of Health Guidelines: D4 for Animal/Arthropod Species: *Drosophila melanogaster* and approved by the Institutional Biosafety Committee (IBS) at UTMB under the Notification of Use (NOU: 2022128)” at the University of Texas Medical Branch (UTMB)

## Credit Authorship Contribution Statement

NM: Data curation, Investigation, Methodology, Writing-original draft; MM: Data curation, Investigation, Methodology, Writing-original draft; AG: Data curation, Writing-original draft; SGS: Data curation, Investigation, Methodology, Writing-review and editing; DCJ: Data curation, Investigation, Software, Writing-review and editing; BK: Conceptualization, Data curation, Funding acquisition, Investigation, Methodology, Project administration, Resources, Software, Supervision, Writing – review and editing

## Declaration of Interests

The authors declare no competing interests.

## Acknowledgements

The authors would also like to acknowledge the Mitchell Center for Neurodegenerative Diseases for the common equipment provision. This work was supported by the Don and Nancy Mafrige Professorship in Neurodegenerative Diseases, Alzheimer’s Association Research Grant (AARG-17-533363 B.K.), National Institutes of Aging (R21-AG059223 and R01-AG063945 B.K.).

## Materials and Methods

### Fly Stocks and Maintenance

All *Drosophila melanogaster* stocks were maintained at 25°C on standard cornmeal-agar-yeast fly food under a 12-hour light:12-hour dark cycle. Specific *Drosophila* strains were procured from Bloomington Drosophila Stock Center (BDSC) at Indiana University: wild-type Canton S (CS) and *w^1118^* mutant lines were kindly provided by Dr. Paul Hardin (Texas A&M University, College Station, TX, USA). The *w^1118^*;+;*UAS-hPLD1* kindly provided by Dr. Raghu Padinjat (National Center for Biological Sciences, TIFR, Bangalore, INDIA). For experiments, newly eclosed flies resulting from genetic crosses were collected into fresh food vials and allowed to acclimatize under standard maintenance conditions for two days prior to electroretinogram (ERG) recording.

### Electrode Preparation

#### Chloritization of Electrical Wires

Electrical wires, serving as the conductive core for the electrodes, were prepared by first carefully removing 6-7 cm of the PVC insulation to expose the inner metal core. Approximately 4-5 such exposed cores were submerged in a 15 mL centrifuge tube containing 100% bleach solution for 24 hours. This chloritization process resulted in the wire cores turning dark brown. Following treatment, the cores were thoroughly rinsed with distilled water, air-dried, and stored for later use in electrode assembly.

#### Fabrication of Glass Micropipettes

For each fly, two glass micropipettes were required: a recording electrode for placement on the eye surface and a reference (grounding) electrode for insertion into the thorax. These micropipettes were fabricated from borosilicate glass capillaries with an outer diameter (O.D.) of 1.5 mm, an inner diameter (I.D.) of 1.10 mm, and a length of 10 cm (BF150-110-10, Sutter Instrument). A Flaming/Brown P-97 micropipette puller (Sutter Instrument) was used with the following optimized parameters: Ramp: 278, Program: 27, Pressure: 500, Heat: 289, Pull: 35, Velocity: 49, and Time: 150.

#### Inspection and Filling of Micropipettes

Pulled micropipettes were carefully stored in a foam-lined petri dish to prevent breakage. Prior to use, each electrode was inspected for cracks or imperfections using a Narishige MF-830 Microforge; any electrode exhibiting damage was discarded and a new one was pulled. Electrodes deemed suitable were back-filled with a 0.9% saline solution using a micro-syringe. After filling, electrodes were visually inspected, often under a dissecting microscope (especially for non-filamentous glass capillaries), to ensure the absence of air bubbles. If bubbles were present, they were dislodged by gently tapping the electrode, allowing them to float to the top and be expelled.

#### Electrode Installation in ERG Apparatus

Filled and inspected micropipettes were then mounted into the ERG recording apparatus. One electrode was positioned in the recording micromanipulator arm and the other in the grounding arm. The chloritized wire within the grounding electrode was connected via an insulated clip to a common metal ground point to ensure proper grounding for the recording setup.

### Fly Mounting and Positioning

#### Anesthetization

Flies were anesthetized using carbon dioxide CO_2_. A CO_2_ gas tank was connected to a *Drosophila* CO_2_ fly pad (Cat# 59-114, Flystuff, Genesee Scientific). Flies within their culture vial were briefly exposed to CO_2_ delivered via a *Drosophila* blow gun (Cat# 54-104, Flystuff) to induce temporary paralysis. Alternatively, we also resorted to cold anesthesia using ice to prevent the long-term effects of CO_2_.

#### Transfer and Preparation for Mounting

Anesthetized flies were gently tapped from the vial onto the CO_2_ conduction plate to maintain anesthesia during the mounting process. It was noted that flies can only remain on the CO_2_ plate for a limited time to avoid asphyxiation, necessitating swift and efficient mounting.

To prepare the mounting apparatus, a standard 10 µL plastic pipette tip was modified. Using a sharp razor blade, the narrow end of the tip was cut approximately 2-3 mm from its extremity, and this small piece was discarded. A second cut was made 4 mm down from this newly cut opening, creating a truncated cone with openings at both ends. Care was taken to ensure the cuts were even to provide a stable surface. This design allowed for the insertion of the fly through the larger opening, with its head and a small portion of the thorax emerging from the smaller, anterior opening.

#### Positioning of Fly in Pipette Tip

Using fine-tipped tweezers, a single anesthetized fly was carefully grasped by its wings and gently inserted, abdomen first, into the larger opening of the prepared pipette tip. The fly was maneuvered such that its head would rest on the plastic lip of the smaller opening, with the back of the head supported by the inner wall of the pipette tip. A wax-tipped needle was then used to delicately push the fly’s body further into the pipette tip until its head and a small portion of the anterior thorax (for grounding electrode placement) protruded from the smaller opening. Extreme care was taken during this step to avoid damaging the fly’s abdomen or thorax. The size of the anterior opening was critical: too small could compress and kill the fly, while too large could allow the fly to escape upon recovery or be pushed back during electrode placement. Some empirical adjustment of the tip dimensions was often necessary.

#### Securing the Mounted Fly

The pipette tip containing the fly was then securely embedded in a small piece of modeling wax adhered to a petri dish. For brief periods, this dish could be inverted onto the CO_2_ plate to maintain anesthesia for approximately 30 seconds longer, preventing premature escape. It was found that orienting the mounted fly at a slight angle within the wax often facilitated easier electrode placement, minimizing the risk of electrode breakage or dislodging the fly. The dish with the mounted fly was then carefully transported to the ERG recording station.

### Electroretinogram (ERG) Recording

#### Equipment Setup and Software

The ERG recording system was powered on sequentially: first the Data Acquisition System (1322 Digidata, Molecular Devices), then the amplifier (Kerr Instruments), and finally the control computer. Once the amplifier indicated it was ready (evia a green indicator light), the gain was set to 250x. The Clampex 8 software (Molecular Devices) was launched to control stimulus delivery and data acquisition.

#### Electrode Placement on Fly

Once the mounted fly was positioned on the ERG stage, its immobility was confirmed. Both the recording and grounding electrodes were initially positioned at a safe distance and height above the fly and then carefully maneuvered towards the fly using micromanipulators, under microscopic observation. The recording electrode was positioned so its tip was slightly above the fly’s compound eye, while the grounding electrode was positioned above the exposed thorax. The microscope zoom was adjusted as needed for precise visualization of the electrode tips relative to the fly.

The grounding electrode was then slowly lowered until its tip pierced the cuticle of the thorax. It was advanced slightly further into the body, observing for any reaction from the fly, ensuring the electrode did not pass through the fly to touch the pipette tip, which could damage the electrode or cause inaccurate readings. If the fly ceased movement or died during this process, it was discarded, and the entire mounting and electrode placement procedure was repeated with a fresh fly. Next, the recording electrode was carefully lowered until its tip made gentle contact with the surface of the eye, causing a slight indentation but without piercing the eye, which would prevent a proper light response.

#### Data Acquisition Protocol

A brief practice light stimulus was often delivered to ensure a proper ERG waveform was being recorded. The experimental protocol typically involved delivering a series of 10 different light impulses (varying in intensity or duration), with each stimulus repeated 10 times, resulting in a total of 100 recordings per fly. After data collection was complete for one fly, it was disposed of, and the process was repeated for subsequent flies.

#### Troubleshooting Notes During Recording

It was noted that glass micropipette electrodes could dull or become clogged between flies, necessitating replacement if the recorded signal amplitude significantly dropped or if electrical noise in the trace increased. In such cases, the chloritization of the internal wires was also checked. If resistance was encountered when attempting to pierce the thorax with the grounding electrode, replacing the electrode tip usually resolved the issue.

### Data Analysis

#### Data Collection and Initial Processing

ERG responses were recorded and saved as .ABF (Axon Binary File) format files. These files were subsequently opened and analyzed using Clampfit 10.7 software (part of pCLAMP, Molecular Devices). The general range of stimulus parameters explored included light pulse durations of 300 ms, 500 ms, 1000 ms, and 2000 ms, and light intensities corresponding to driver currents of 5 pA, 10 pA, 15 pA, and 20 pA, applied in various combinations according to specific experimental protocols.

#### Baseline Adjustment and Noise Filtering

Raw ERG traces were first adjusted to a common baseline to allow for accurate amplitude measurements across different recordings and genotypes. This approach was consistently applied to all genotypes. The recordings sometimes exhibited slight electrical interference noise, characterized by minute, constant oscillations (visible in Figure 1 panel A of the referenced figures). This noise was minimized post-acquisition using the “Electrical Interference Filter” feature in the analysis software to remove external harmonics, resulting in a cleaner trace (as shown in panel B of the referenced figures). The nadir (peak negative deflection) and peak positive deflection of the ERG waveform were marked using cursors to determine the amplitudes of different components. For quantitative analysis, the values from 10 repeated traces for each stimulus condition were typically averaged.

## Supplementary Information

### MATERIALS

| **REAGENTS** | **DESCRIPTION** | **COMPANY &**<br>**CATALOGUE #** |
| --- | --- | --- |
| 100% Bleach Solution | Decontaminant (add catalogue number) | Clorox: USA001229 |
| Fly Stock 33799 | RRID:BDSC_33799 | Bloomington *Drosophila* Stock Center |
| Fly Stock 458 | RRID:BDSC_50892 | Bloomington *Drosophila* Stock Center |
| Fly Stock 32040 | RRID:BDSC_32040 | Bloomington *Drosophila* Stock Center |

### EQUIPMENT

| **NUMBER** | **EQUIPMENT** | **COMPANY &**<br>**CATALOGUE #** |
| --- | --- | --- |
| 1 | Flaming/Brown Micropipette Puller | Sutter Instruments<br>Company: P-97 |
| 2 | W.F. 15X Eye piece | Narishige: MF-830<br>MicroForge |
| 3 | 15mL High-Clarity Polypropylene<br>Conical Tube | Falcon: 14-432-22 |
| 4 | 50mL Conical Centrifuge Tubes | ThermoFisher Scientific:<br>339650 |
| 5 | Extracellular Recording Amplifier | Kerr Lab: 8546850 |
| 6 | Fire polished Borosilicate Glass with<br>Filament | Sutter Instruments<br>Company: BF150-110-10 |
| 7 | ACE 1 Fiber Optic Light Source | Schott: A20500 |
| 8 | Flowbuddy Flow Regulator | Flystuff: 59-122B |
| 9 | Stereo Microscope | Olympus: SZ51 |
| 10 | Hook Up Wire, Solid, Type 18 Gauge,<br>100' Length, 300V, Red | NTE Electronics: WHS18-02-100 |
| 11 | Disinfecting Bleach | Clorox: USA001229 |
| 12 | Stereomicroscope | Wild: M5 |
| 13 | Magnet Holder Base | Kanetec: MB-PB |
| 14 | S2 Amplifier | Kerr Scientific: 3000-521 |
| 15 | 16-Bit Acquisition System | Digidata 1322A Molecular<br>Devices / Axon<br>Instruments: 83029-1 |
| 16 | Red film | Pangda: 4331997009 |
| 17 | Aluminum Foil Roll | Fisherbrand: 01-213-100 |
| 18 | Carbon Dioxide USP 50 LB | UTMB LINDE: CD M-50 |

### GENERAL SUPPLIES

| **NUMBER** | **SUPPLIES** |
| --- | --- |
| 1 | Paper Towels |
| 2 | Kimwipes |
| 3 | Gloves |
| 4 | Pens |
| 5 | Timer |
| 6 | Permanent Marker |
| 7 | Forceps |
| 8 | Scissors |
| 9 | Paint Brush |
| 10 | Fly Vials |
| 11 | Vial Plugs |

### *DROSOPHILA* FLY - FOOD MAKING PROTOCOL

|  |  |
| --- | --- |
| **INGREDIENTS - 3L OF** | **METHOD** |
| **FOOD** |  |
| Sugar - 37.05 g | 1. Added first 3 ingredients to 3000 mL of dH<sub>2</sub>O in mixing bowl |
| Yeast - 37.05 g | 2. Mixer to optimal speed (much less than 5-10 mins to boil) |
| Jazz Mix - 467.22 g | 3. Turned on heat. Heated to boiling, then shut off heat for the remainder of the protocol |
| Propionic Acid - About 24 mL | 4. When food cooled to ~75° C, propionic acid was slowly added |
| dH<sub>2</sub>O | 5. When food cooled to ~67° C, food was poured into vials |
|  | 6. 4 trays were immediately plugged and left at room temperature to set overnight |
|  | 7. Trays set well. One tray was placed in fridge and the other 3 trays were placed into the -20° C freezer |

### Methods

#### Preparation of Flies

All *Drosophila* stocks were maintained at 25°C on standard fly food under a 12:12-h light-dark cycle. Drosophila stocks were either purchased from Bloomington Drosophila Stock Center (BDSC), Bloomington, Indiana, USA or provided. Flies were collected and acclimatized for 2 days.

#### Preparation of Electrodes

Electrical wires were prepared by removing insulation and chloritizing in 100% bleach for 24 h. Two electrodes (recording and grounding) were required per fly. Borosilicate glass microelectrodes (O.D.: 1.5mm, I.D.: 1.10mm) were pulled using a Flaming/Brown puller with specific settings (Ramp: 278, Program: 27, Pressure: 500, Heat: 289, Pull: 35, Velocity: 49, Time: 150). Electrodes were inspected using a microforge and replaced if cracked. They were filled with 0.9% saline, checked for bubbles, and placed in the ERG apparatus. The grounding wire was connected to an insulated clip.

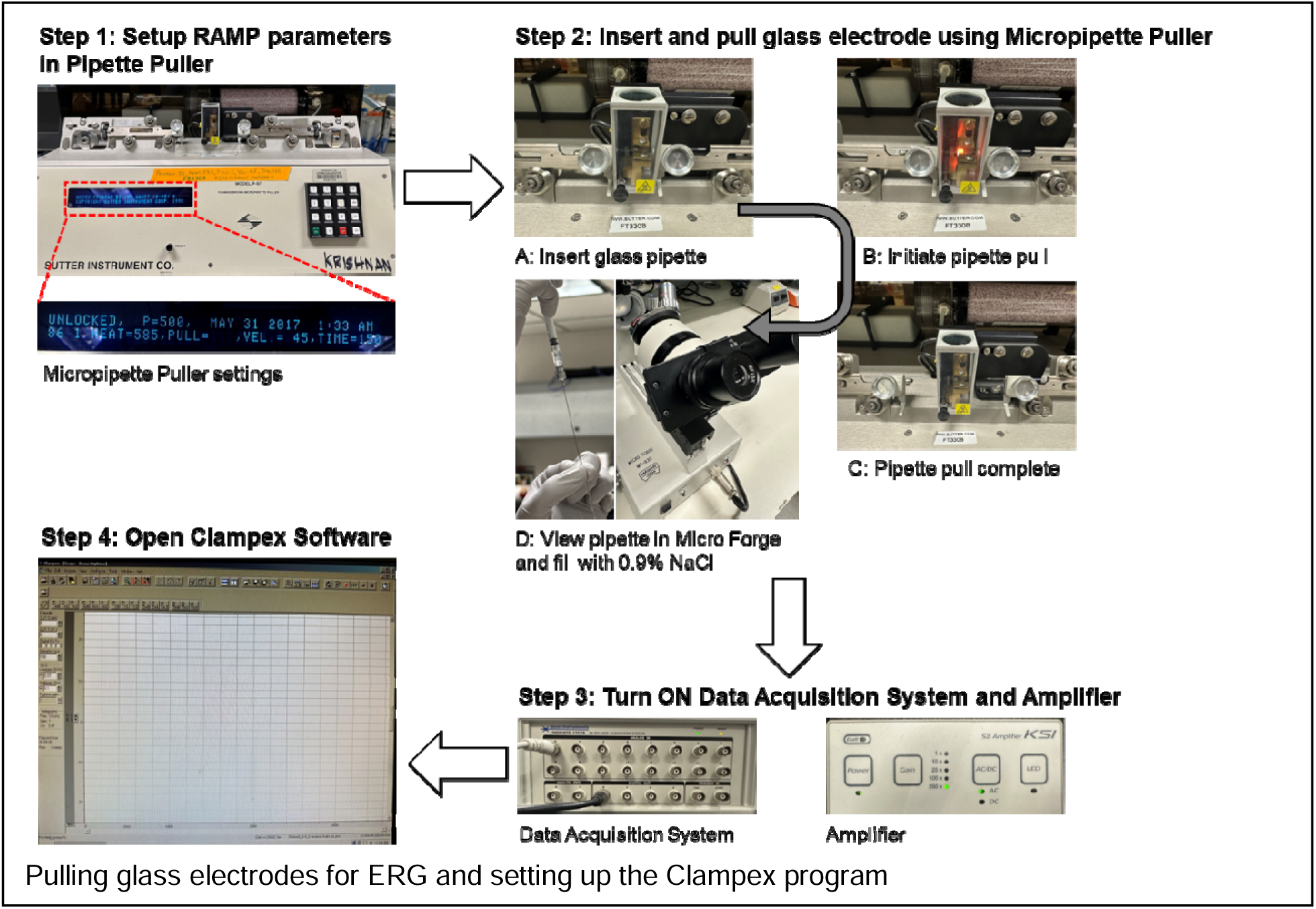

#### Placing Flies in Electrode

Flies were anesthetized using CO_2_. A 10 µL pipette tip was cut to create a holder (4mm length). A fly was inserted using tweezers and gently pushed with a wax-tipped needle until its head and part of the thorax protruded. The tip was secured in wax on a dish, often at an angle.

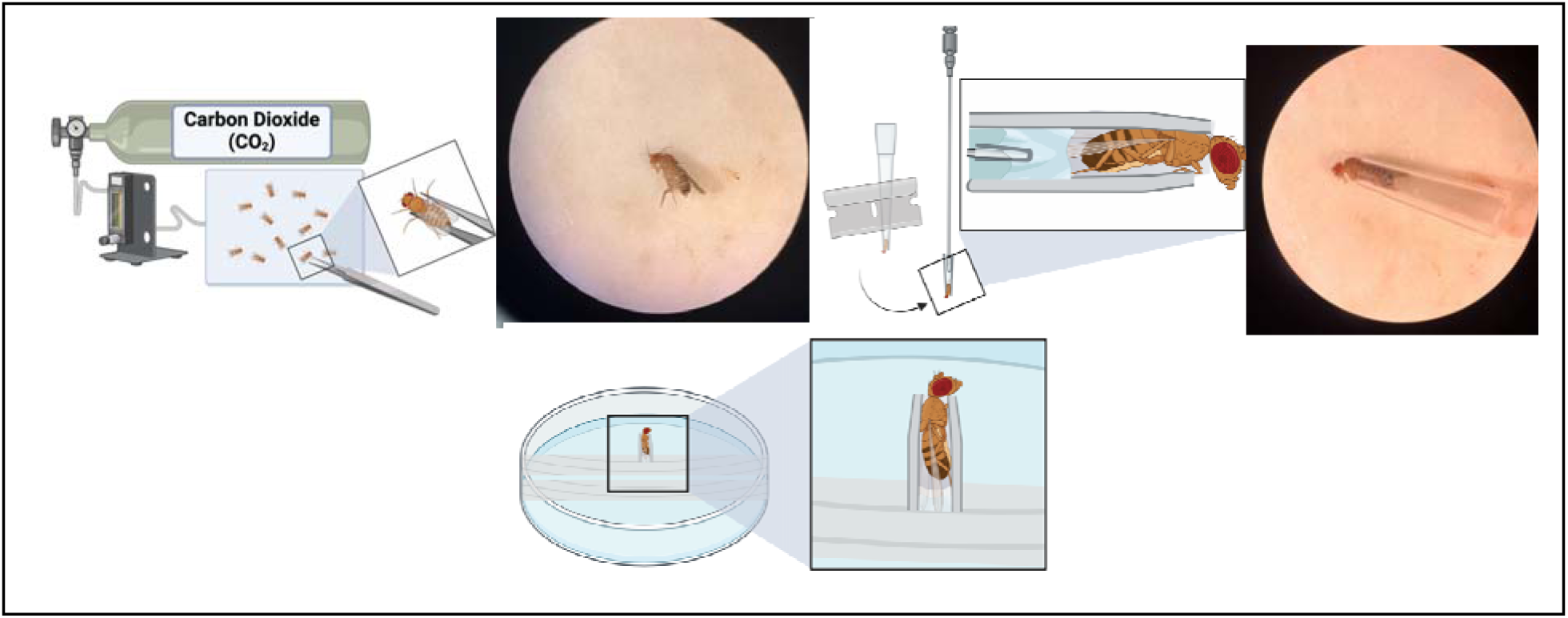

#### ERG Recording

The Data Acquisition System, Amplifier (gain set to 250x), and Clampex 8 software were started. Electrodes were positioned carefully: grounding electrode into the thorax and recording electrode indenting (but not piercing) the eye. Recordings were taken using a program with 10 different light impulses, repeated 10 times. Electrodes were replaced if readings dropped or noise increased.

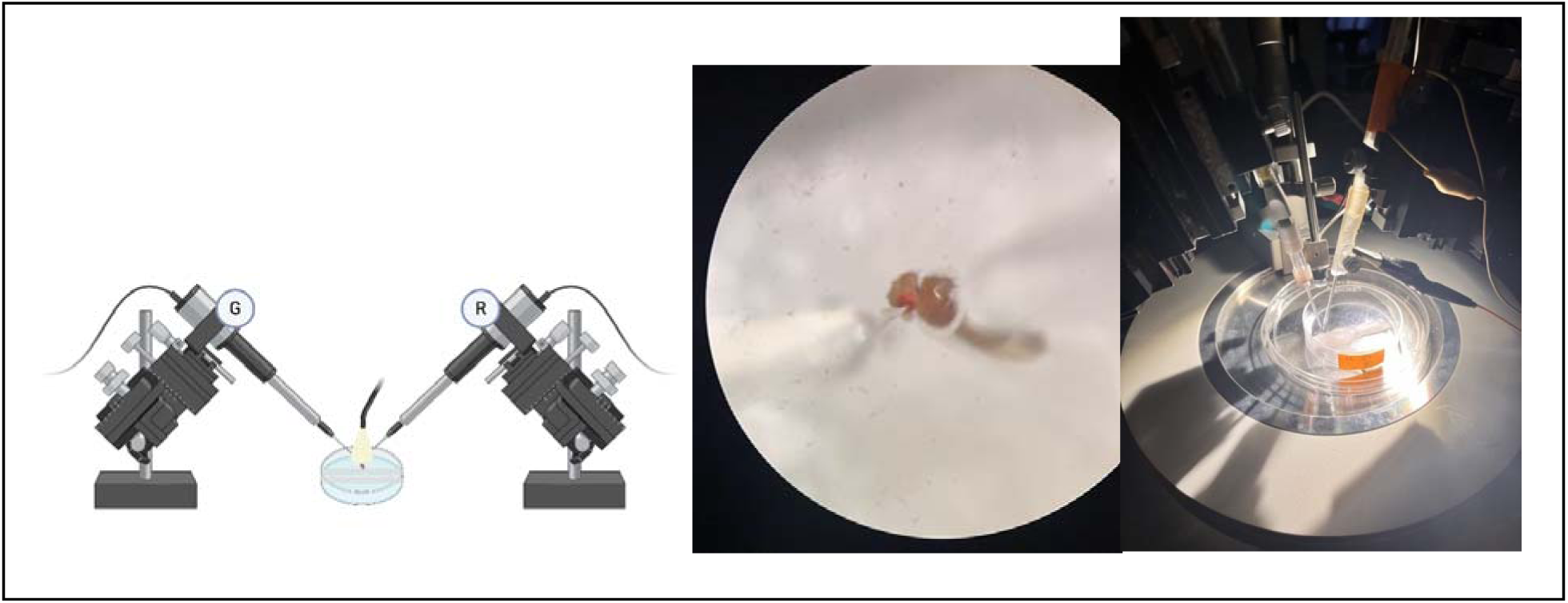

#### Data Analysis

Data was collected in ABF files and analyzed with Clampfit 10.7. Raw data was baseline-adjusted. Electrical interference was minimized using a filter. Amplitudes were determined, averaged, and used for analysis. A two-way ANOVA was used for statistical comparison.

## Notes

### Competing Interest Statement

The authors have declared no competing interest.

### Summary of Updates

Abstract updated to align more with paper's focus. Revisions made across sections to incorporate new data

